# Humanized mice enable in vivo evaluation of engineered plasma cell biology and therapeutic function

**DOI:** 10.1101/2025.07.14.664835

**Authors:** Tyler F. Hill, Anna E. Helmers, Rene Yu-Hong Cheng, Swati Singh, Leah J. Homad, Parnal Narvekar, Gregory D. Asher, Nathan D. Camp, Emma R. Suchland, Andee R Ott, Malika Hale, Chris D Thouvenel, Claire M. Stoffers, Stefan Lachkar, Andrew T. McGuire, David J. Rawlings, Richard G. James

**Affiliations:** University of Washington, Medical Scientist Training Program, Seattle WA; Seattle Children’s Research Institute, Center for Immunity and Immunotherapy, Seattle WA; Fred Hutchinson Cancer Center,, Vaccine and Infectious Disease Division, Seattle; University of Washington, Departments of Global Health and Lab Medicine and Pathology,, Seattle WA; University of Washington, Departments of Pediatrics and Immunology, Seattle WA; University of Washington, Departments of Pediatrics and Pharmacology, Seattle WA

## Abstract

Engineered plasma cells (ePCs) offer a durable strategy for in vivo delivery of therapeutic antibodies, but standard immunodeficient mouse models lack human immune factors critical for plasma cell survival and function. We utilized a humanized mouse model in which NOD.Cg-*Prkdc*^*scid*^ *Il2rg*^*tm1Wjl*^/SzJ (NSG) mice were engrafted with human CD34^+^ stem cells as recipients for infusions with autologous ePCs. In this setting, ePCs localized to plasma cell niches and stably secreted antibodies for over three months. To improve the selection of antibodies for secretion, we developed a B cell receptor surface display screen that identified candidate antibody sequences with high secretion potential. An anti-SARS-CoV-2 antibody (clone 297) selected by this method showed robust secretion both in vitro and vivo, and serum from ePC-engrafted mice potently neutralized SARS-CoV-2 pseudovirus. Together, these findings establish a physiologically relevant model for testing human ePCs, and offer a generalizable strategy for optimizing antibody selection to support long-term therapeutic delivery.

## Introduction

Diseases treated through repeat infusions or injections of recombinant proteins may benefit from continuous, *in vivo* therapeutic delivery. For example, patients receiving antibody therapies for autoimmune disorders, cancers or infectious disease often require regular infusions to maintain clinical benefit, which can pose logistical and quality of life challenges^1–3^ that limit real world effectiveness. To address these challenges, several groups have developed gene^4–6^ and cell therapy-based^7^ approaches to provide sustained in vivo production of therapeutic antibodies, with the goals of improving pharmacokinetics, and reducing treatment burden.

Among these strategies, engineered B cells,^8–11^ and their differentiated counterparts, plasma cells (PCs)^12,13^ offer a promising platform for continuous antibody secretion. Human PCs naturally possess both a long lifespan^14^ (estimated half-life of 11 to 200 years^15^) and extraordinary secretory capacity (up to 10,000 IgG molecules per second),^16–18^ making them attractive for long-term protein delivery. Engineered PCs (ePCs) derived ex vivo from B cells have been shown to resemble endogenous human PCs in phenotype and function,^19^ and have successfully delivered therapeutic antibodies in preclinical models, which include those targeting viral antigens,^12^ leukemia-associated surface receptors,^20^ and immune checkpoints.^13^ Finally, early-phase clinical trials (NCT05682144) using ePCs to produce an enzyme replacement for mucopolysaccharidosis type I,^21^ have demonstrated promising safety profiles and potential functional improvement. Collectively, these advancements underscore the therapeutic potential of ePCs and the importance of developing robust in vivo models to study their behavior, longevity, and efficacy within a humanized immune system context.

Most studies to date have assessed ePC persistence and function using adoptive transfer into immunodeficient mice strains such as NOD.Cg-*Prkdc^scid^ Il2rg^tm1Wjl^*/SzJ (NSG) mice.^19,22^ However, these models lack many components of the human immune system that influence PC survival, homing and function. NSG mice reconstituted with human CD34+ stem cells (NSG-huCD34) offer a more physiologic environment for ePC studies,^23^ as they harbor multiple human immune lineages, including CD4^+^ T cells,^24^ and macrophages,^25^ which are critical for PC maintenance. Despite this, endogenous PCs in NSG-huCD34 mice typically remain IgM-restricted due to poor class-switching in the absence of inflammatory cues and efficient T cell help.^26^ This feature creates a unique advantage for ePC studies, as it facilitates tracking of engineered IgG secreting cells by total and antigen-specific IgG titers.

Taken altogether, these features make the NSG-huCD34 model well suited for evaluating the persistence, localization, and function of IgG ePCs in vivo. Here, we demonstrate that the engraftment of ePCs was more efficient in mice engrafted with syngeneic CD34 cells and resulted in significantly higher and more stable levels of human endogenous and antigen-specific antibodies. To boost anti-pathogen antibody levels, we developed a surface BCR screen to find antibody sequences with improved secretion capacity. Finally, sera from humanized mice engrafted with ePCs secreting anti-SARS-CoV-2 antibodies were capable of neutralizing SARs-CoV-2 pseudovirus. This study provides proof of concept that the syngeneic ePC NSG-huCD34 mouse model can be used as therapeutic ePC testing platform and suggests that ePCs have the capacity to provide long lasting therapeutically functional antibodies.

## Results

### Humanized mice enhance the engraftment and longevity of human PCs

We first wanted to establish whether mice humanized with hematopoietic stem and progenitor cells produce human factors capable of supporting engraftment and longevity of human PCs.^19,22,27–32^ To test this, we implanted NOD.Cg-Prkdc^scid^ Il2rg^tm1Wjl^/SzJ (NSG) mice with CD34^+^ hematopoietic stem and progenitor cells purified from healthy donor peripheral blood mononuclear cells (PBMCs). Ten to twelve weeks after establishment of the human hematopoietic system, we sacrificed the mice and analyzed tissues and serum for evidence of supportive factors and cell types.^24^ Spleens harvested from humanized mice were larger than those from non-humanized controls and contained ordered sections of white pulp (lymphoid compartment; Figure S1A). As previously shown by Shultz *et al.*,^33^ the spleens and bone marrow from NSG-huCD34 mice contained human myeloid cells, B cells, and, to a lesser extent, CD4^+^ T cells (Figure S1B). Serum from these mice contained measurable levels of human B-cell activating factor (BAFF), interleukin-6 (IL-6) and interleukin-21 (IL21), which was not detected in control NSG mice (Figure 1A), along with other human cytokines (Figure S1C). These findings indicate that NSG-huCD34 mice recapitulate key human environmental features needed to support engraftment and persistence of long-lived human PCs, surpassing prior immunodeficient models in this regard.

**Figure 1:**
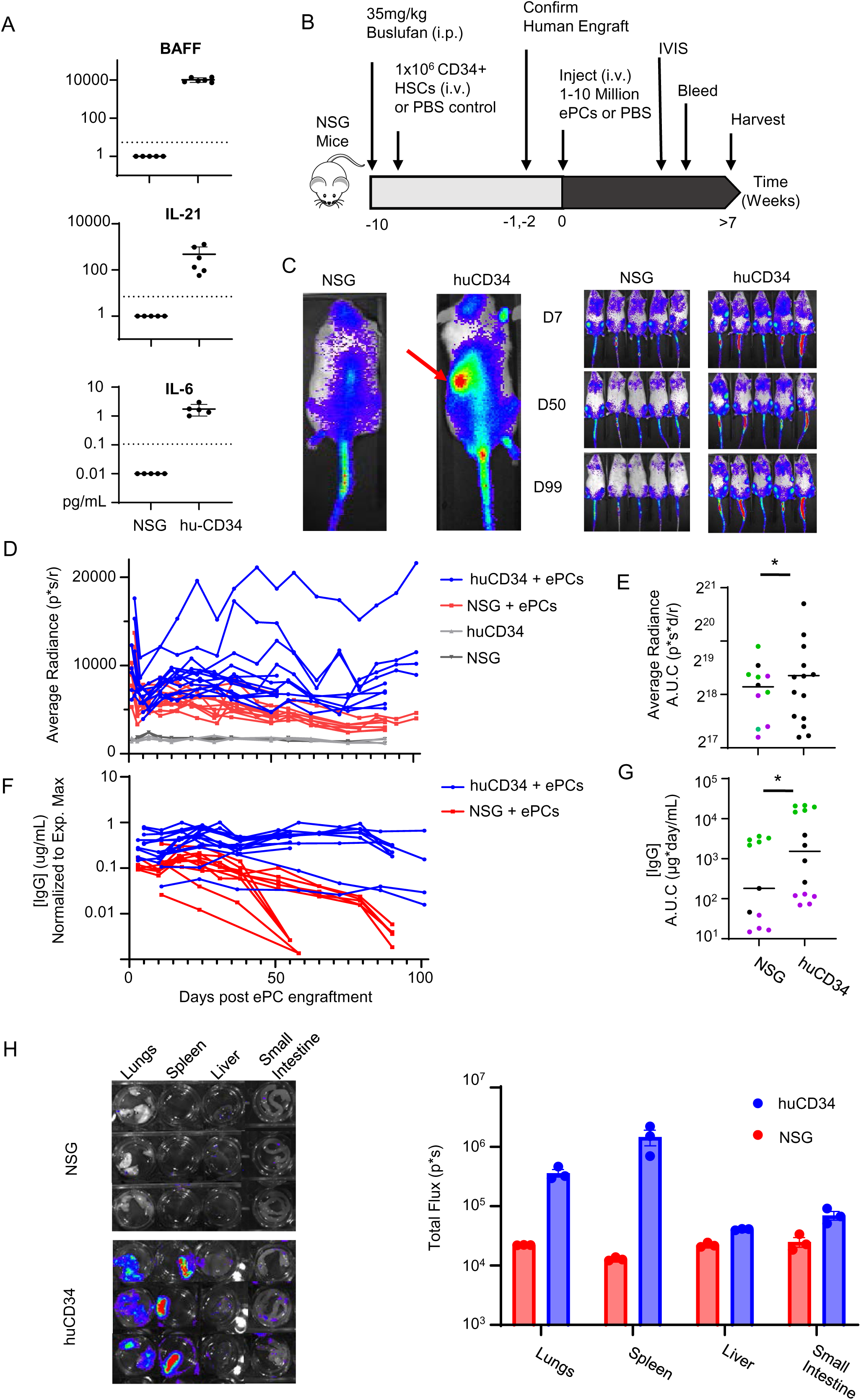
Hematopoietic stem and progenitor cell humanization of NSG mice permits robust engraftment of human ePCs. Immunodeficient NSG mice were injected with huCD34^+^ hematopoietic stem and progenitor cells via tail vein injection. Mouse tissues and peripheral sera were harvested 10 weeks after engraftment. **A)** Cytokine concentrations in the sera of NSG-huCD34 and NSG mice were measured by ELISA. Dotted line indicates lower limit of detection. **B)** Experimental schema for the assessment of luciferase expressing ePCs engrafted in NSG and NSG-huCD34 mice. Mice were injected (s.c.) with luciferin and bioluminescence emitted from luciferase expressing ePCs was measured at various time points. **C)** Representative bioluminescence images of NSG-huCD34 and NSG mice 21 days after engraftment with ePC shown in the left panel. Red arrow emphasizes additional engraftment in the spleen of a representative humanized mouse. Additional representative bioluminescence ventral images of NSG and NSG-huCD34 mice at various times after ePC are shown in the right panel (color scale; min: 5 x 10^3^; max: 1 x 10^5^). **D)** Bioluminescent average radiance was quantified from each mouse over time. **E)** Area under the curve analysis of average radiance was conducted. Different colors correlate to three independent experiments with three different donors. **F)** The concentration of human IgG was measured via ELISA of peripheral sera and normalized to the maximum concentration achieved in each experiment. **G)** Area under the curve analysis of the IgG titers was conducted. Different colors correlate to three independent experiments with three different donors. **H)** Mice 21 days after engraftment with ePCs were perfused with PBS containing luciferin and tissues were harvested for *ex vivo* bioluminescence imaging (color scale; min:4×10^3^ max:1×10^5^). Luminescence from each sample was measured. A) Data across 2 donors. D-G) Data across 3 donors in 3 independent experiments with p-value calculated by unpaired student’s t test (* p < .05). H) Data from one donor.

We engineered syngeneic PCs to assess the engraftment kinetics of ePCs in NSG-huCD34 mice. We began with G-CSF mobilized peripheral blood mononuclear cells (PBMCs) that were collected and cryopreserved at the time of the CD34^+^ hematopoietic stem and progenitor cell isolation. Using methods we have previously described for non-mobilized PBMCs^19^ (schematized in Figure S2A), we employed homology-directed repair to introduce either blue fluorescent protein (BFP) or *firefly* luciferase gene cassettes into purified B cell populations. These engineered B cells were differentiated into ePCs, and evaluated for gene integration (Figure S2B), reporter expression (Figure S2C), and immunophenotype (Figure S2D). Consistent with our prior findings,^19,22^ greater than 20% of the differentiated mobilized cells expressed human PC markers (CD38^+^CD138^+^, Figure S2D) and efficiently produced class-switched Immunoglobulin G (IgG) ePCs (Figure S2E). These results show that syngeneic trackable human ePCs can be generated from G-CSF-mobilized PBMCs left over from CD34^+^ isolations.

To determine whether humanized mice can more efficiently support PC engraftment and maintenance, we adoptively transferred luciferase-expressing ePCs into syngeneic NSG-huCD34 mice (Figure 1B). Similar to non-humanized control animals, bioluminescent ePCs were detected in the bones, tail and lung in NSG-huCD34 mice (Figure 1C). However, NSG-huCD34 mice also exhibited signals in the spleen that were not evident in non-humanized controls (arrow, Figure 1C). Furthermore, over time the luminescence signal was increased in the NSG-huCD34, relative to the NSG control mice (Figure 1D-E and Figure S3A). Consistent with the bioluminescence data, we found that human IgG, which is entirely derived from the ePCs in the huCD34 model,^33,34^ remains stable in the NSG-huCD34 for the duration of the study (∼100 days), but not NSG recipients (Figure 1F-G, Figure S3B). We calculated the cellular half-life of ePCs using bioluminescence decay after day 30 (Figure S3C), and found that the ePC half-life is substantially greater in NSG-huCD34 mice (∼238 days) relative to non-humanized controls (∼58 days).

Lastly, to determine if humanized support was present in tissues not readily evaluated using whole animal luminescence, we prepared the lungs, spleen, liver, and small intestine and imaged them *ex vivo* twenty days after ePCs engraftment. We detected a trend towards increased luminescence in all tissues from NSG-huCD34 mice, which was most pronounced in the lungs and spleen (Figure 1H). Collectively, these findings show that partial reconstitution of a human hematopoietic system improves the engraftment and persistence of ePCs in several organs.

### PCs are retained in the bone marrow and spleen in NSG-huCD34 mice

Next, we determined whether the spleen-or bone marrow-localized ePCs exhibited canonical PC phenotypes in NSG-huCD34 mice. To do this, we collected tissues for immunohistology from mice engrafted with luciferase-edited ePCs and isolated cells for flow cytometry from mice engrafted with BFP-edited ePCs. As expected, human PCs (huCD3^-^ CD38^+^huCD45^+^CD138^+^) were detected in the bone marrow and spleens of NSG-huCD34 mice regardless of ePC administration (Figure 2A-B, additional flow gating Figure S4A). However, only mice that received ePCs harbored class-switched (IgG^+^) PCs in these tissues, whereas control NSG-huCD34 mice had only IgM^+^ PCs (Figure 2B). Importantly, we were able to find reporter-positive PCs (BFP^+^CD38^+^ CD138^++^) in the spleens and bone marrow of mice receiving BFP-expressing ePCs (Figure 2C, Figure S4B). Using immunohistochemistry, we found that all luciferase positive cells in the bone marrow (femur; Figure S4C) and spleens (Figure 2D) of NSG-huCD34 mice co-expressed CD138. These findings demonstrate that ePCs migrate to bone marrow and spleen niches of NSG-huCD34 mice, where they can retain PC phenotypes and persist over time.

**Figure 2:**
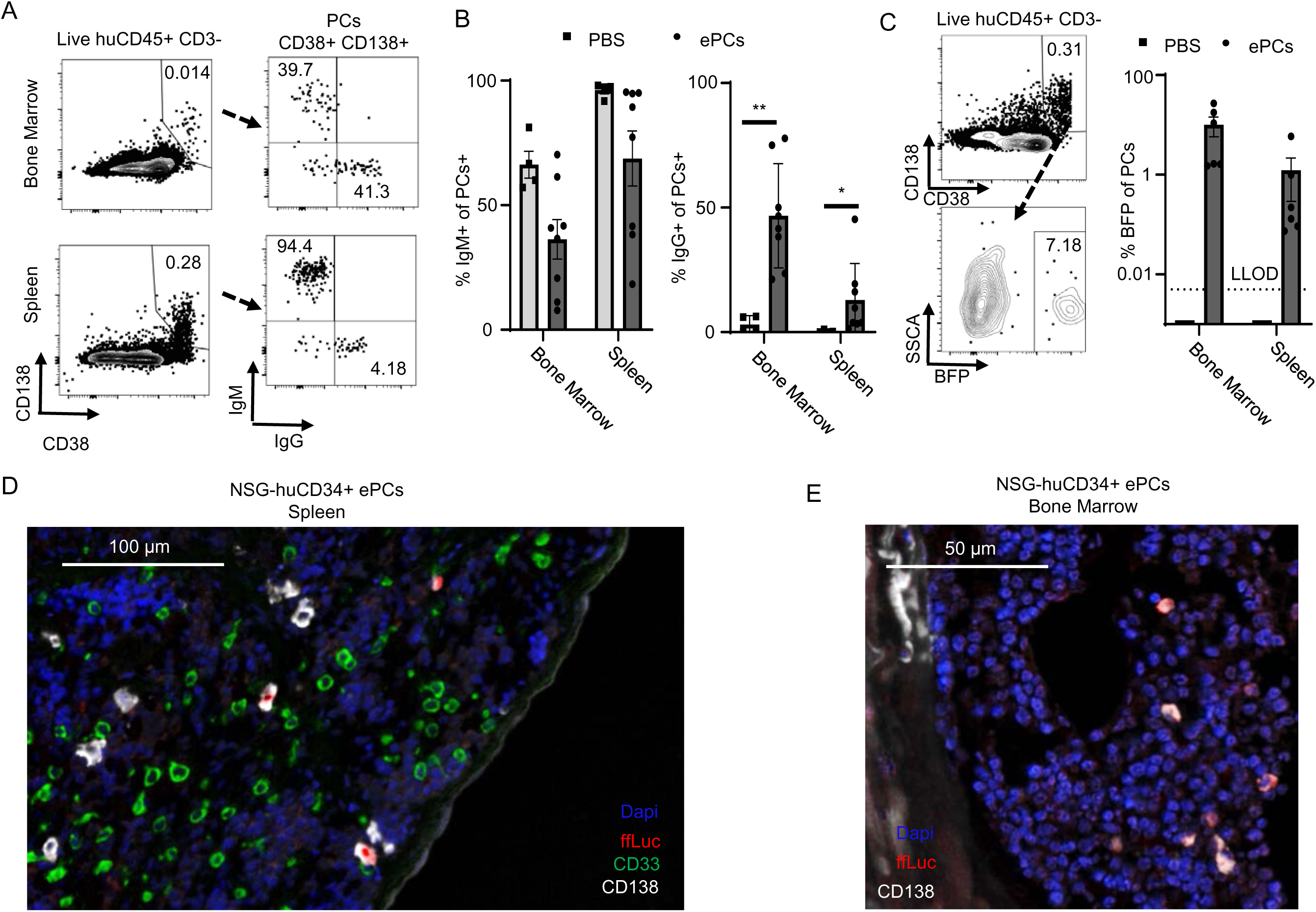
ePCs maintain PC phenotypes and persist in the bone marrow and spleens of humanized mice. Mouse bone marrow and spleens were harvested between 5 and 10 weeks post ePC engraftment in NSG-huCD34 mice. **A)** Representative flow plots of an ePC engrafted NSG-huCD34 mouse shows the staining and gating strategy used to determine the isotype of PCs found in the bone marrow and spleens of ePC engrafted NSG-huCD34 mice. **B)** The percentage of PCs (huCD45^+^,CD38^+^,CD138^+^) that express IgM and IgG were quantified and graphed, revealing that only mice receiving ePC have IgG^+^ PCs. **C)** Representative flow plots shown in left panel and quantification of the percent of PCs that express BFP shown in right panel. **D)** Immunohistology of a representative spleen from ePC engrafted NSG-huCD34 mice showing colocalization of Luciferase with CD138 positive cells. Spleens were also stained for human CD33 and DAPI. B-C) Data across two donors in two independent experiments with P value calculated by unpaired student’s t test (** p < .01; * p < .05).

### PCs engineered to secrete engineered antibodies exhibit longevity in humanized mice

Next we tested whether ePCs expressing therapeutic antibodies exhibit similar engraftment and antibody production kinetics as unedited PCs in NSG-huCD34 recipients. Using an approach previously described in Moffett et al^12^ (Figure 3A), we introduced a single-chain version of the anti-Epstein Barr Virus (EBV) antibody, AMMO1.^35^ To enable tracking of AMMO1-expressing ePCs, we incorporated a Strep-Tag II (STII) linker peptide between the light and heavy chain sequences. Following gene editing and differentiating B cells into ePCs (Figure S5A), we observed intracellular STII staining at rates comparable to those achieved with earlier reporter templates (Figure 3B). Moreover, AMMO1-specific IgG was detectable in the culture supernatants by ELISA using EBV-specific gHgL^35^ as the capture antigen (Figure S5B-C).

**Figure 3:**
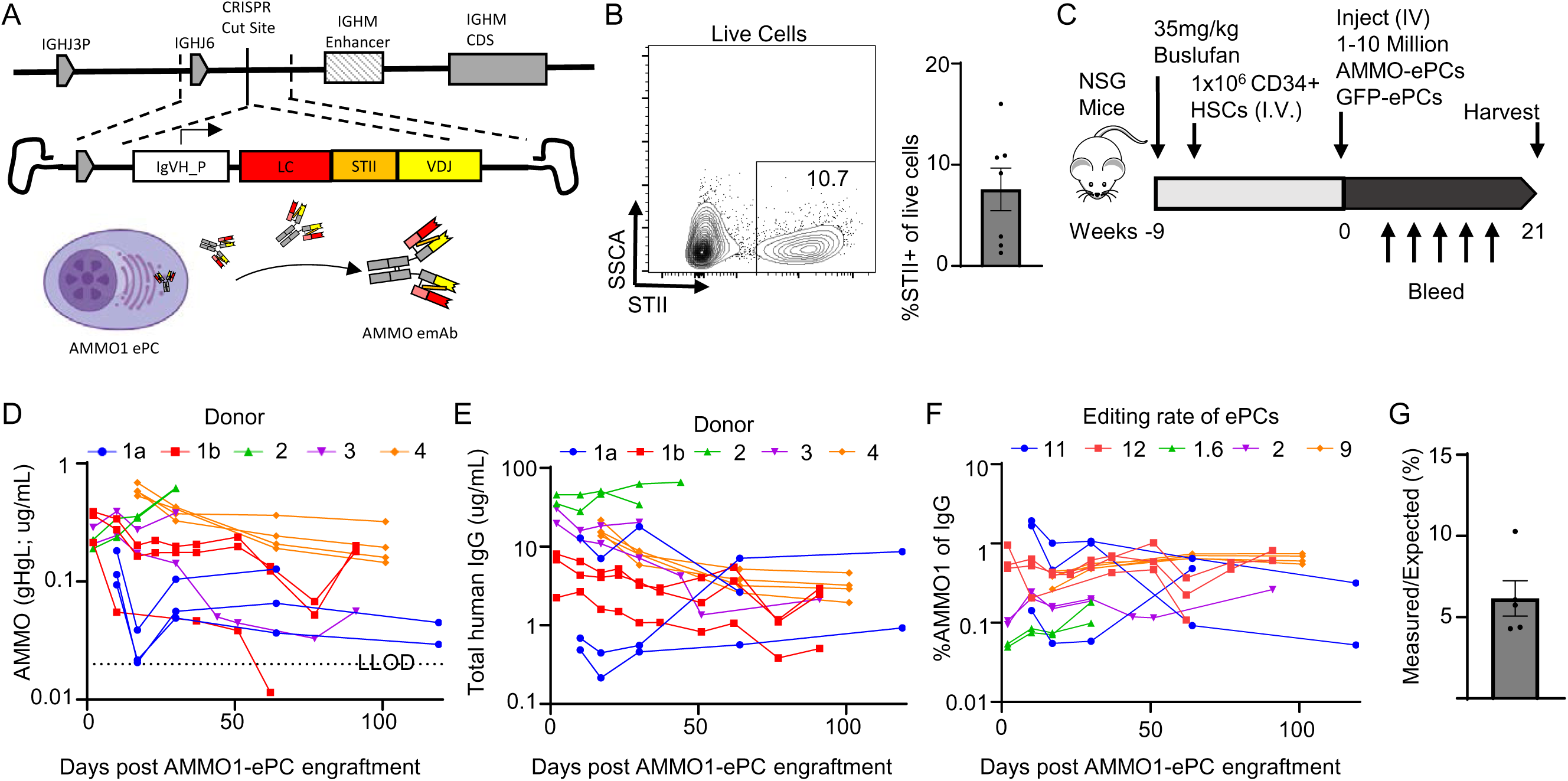
ePCs persist and stably produce engineered antibodies in NSG-huCD34 mice. **A)** Schematic showing the genome editing strategy used to generate ePCs expressing AMMO1. **B)** 5 days after gene editing, ePCs were intracellularly stained for the STII epitope tag and analyzed by flow cytometry. **C)** Experimental timeline for the assessment of AMMO-ePC engraftment in NSG-huCD34 mice. **D)** Secreted AMMO1 IgG and **E)** total IgG was quantified in the sera from mice engrafted with ePCs using antigen specific (gHgL) or IgG ELISA. **F)** AMMO1 IgG levels were compared to total IgG levels and graphed. **G)** Percent of expected AMMO1 concentration calculated based on AMMO1 editing rates and the ELISA values; these were plotted. B-E) Data across 3 donors in 2 independent experiments. G-H) Data across 4 donors in 5 independent experiments.

To assess whether ePCs secreting AMMO1 IgG exhibited long-term maintenance in animals, we next infused AMMO1- or GFP-expressing ePCs into NSG-huCD34 mice (see schematic, Figure 3C). AMMO1 was reproducibly detected in serum from mice receiving AMMO1 ePCs across five experiments, though abundance varied by donor (Figure 3D). In contrast, AMMO1 IgG was undetectable in all mice receiving GFP ePCs. Following infusion, AMMO1-IgG abundance remained stable in NSG-huCD34 serum for up to 100 days (Figure 3D). As we previously reported, total IgG levels in these mice were also stable over the duration of the experiment (Figure 3E). Importantly, when normalized to total IgG, we found the percentage of AMMO1 correlated with the editing rate in each experiment and remained constant over time (Figure 3F), suggesting that AMMO1-expressing cells did not suffer a selective engraftment or maintenance disadvantage relative to unmodified PCs within the same infusion product.

However, the observed serum concentration of AMMO1-IgG was approximately five percent (CV:39.61%) of the expected value based on the frequency of edited cells, and total IgG abundance (Figure 3G). To rule out affinity as a limiting factor, we confirmed that the recombinant, single-chain AMMO1 antibody exhibited binding affinities similar to the split chain version (Figure S5B), suggesting that reduced detection by antigen-specific ELISA was not due to impaired target binding. Re-analysis of *in vitro* supernatant ELISAs for AMMO1 (Figure 5SC) and total IgG (Figure 5SD), in conjunction with editing rates (Figure S5E), revealed that AMMO1 secretion *in vitro* was reduced to a similar extent (to ∼5%) as observed *in vivo* (Figure S5F).

**Figure 4:**
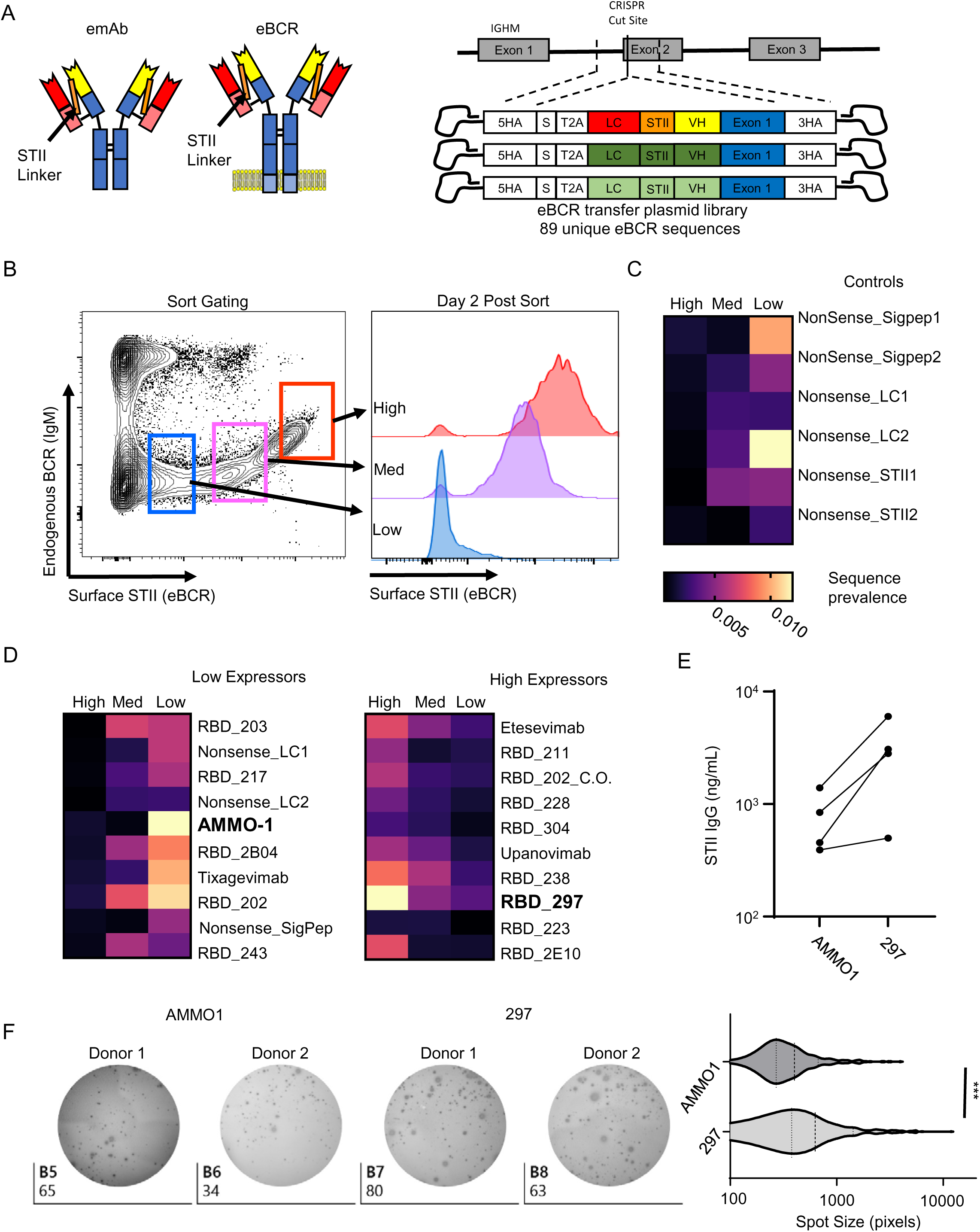
eBCR screen identifies antibodies capable of high secretion from ePCs. **A)** Representation of BCR transfer plasmid library and genome editing strategy. **B)** eBCR library engineered Ly-7 cells were stained for surface endogenous BCR (IgM) and for STII. The stained cells were then sorted according to the gates shown (left panel). To confirm stability of the edited population, each sorted population was re-stained for STII two days post-editing (right panel). Genomic DNA was then isolated from each sorted population and the eBCR frequencies were quantified by deep sequencing. **C)** Sequence frequency for each nonsense control was mapped by a heat map for each sorted population. D) Heat maps showing the relative sequence frequency of the 10 least and most expressed eBCRs. E) The “high expression” BCR sequence for 297 was used to generate 297 ePCs. Secreted antibodies were measured from supernatants from AMMO1- and 297-ePCs using anti-STII ELISA. Data generated using four donors across two independent experiments. F) Anti-STII ELISpot was performed on Day 10 ePCs and spot size was measured. Data was generated using two donors with P value calculated by paired student’s t test (*** p < .001).

**Figure 5:**
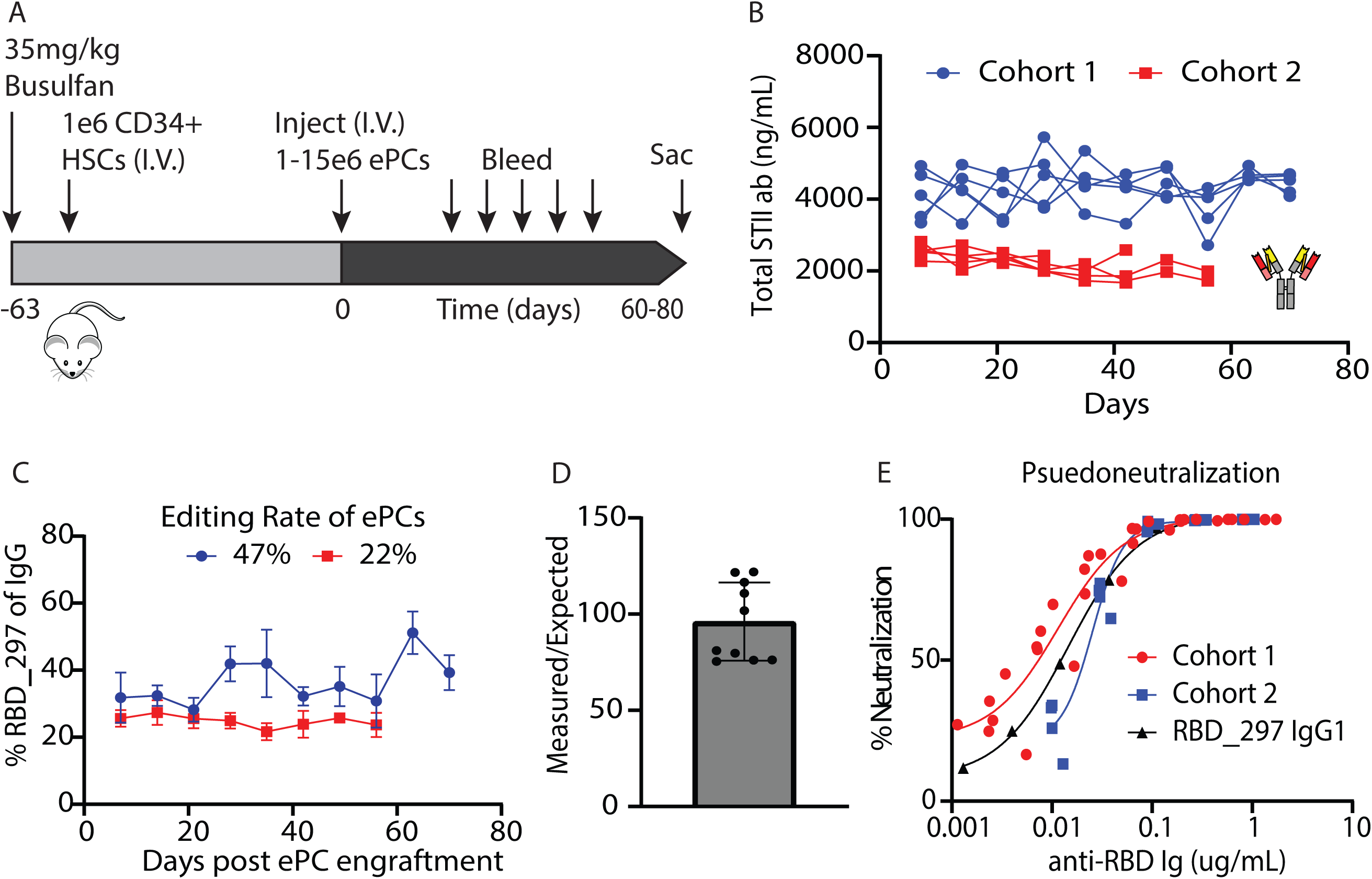
Sera from humanized mice engrafted with ePC is capable of neutralizing SARS-CoV-2 pseudovirus. **A)** Experimental timeline for the assessment of 297-ePC engraftment into NSG-huCD34 mice. **B)** Secreted 297 antibodies in the sera of ePC engrafted mice was measured by STII-ELISA. **C)** 297 IgG levels were compared to total IgG levels. **D)** Average measured 297 antibody concentration was compared to the expected 297 concentration, as determined by 297 editing rates measured by flow cytometry. **E)** Sera was incubated with SARS-CoV-2 pseudovirus containing a luciferase transgene and applied to ACE2-expressing 293Ts. Two days later, viral infection was measured by luminescence. Luminescence values were normalized to those measured in NSG control sera and graphed as percent neutralized. **B-D)** Data across one donor in two independent experiments.

While the amount of AMMO1 in serum was insufficient to neutralize EBV pseudo-virus and secretion was lower than predicted, these findings demonstrate that monoclonal antibody producing ePCs can stably engraft in humanized mice (>90 days), without a selective disadvantage due *in vitro* engineering.

### Surface display screen identifies highly secreted engineered monoclonal antibodies

Previous reports have also shown that engineered BCR surface expression varies considerably depending on the antibody variable domain sequence.^10,12^ We hypothesized that these differences in BCR expression would correlate with antibody secretion levels from ePCs. To test this, we leveraged the human lymphoma cell line (OCI-Ly7^36^), which is dependent on surface BCR, to evaluate BCR surface expression across a library of antibody variable domain sequences. We designed an engineering strategy targeting the second exon of IGHM (Figure 4A) using an sgRNA that disrupts endogenous BCR expression in the absence of homology-directed repair,^37^ thereby enabling enrichment of successfully edited cells. We then constructed a repair template library consisting of 79 unique antibody variable domain sequences (Supplemental Table 1), including clones specific to diverse antigens such as EBV and the receptor binding domain of the SARS-CoV-2 spike protein. Control sequences with altered codon optimization schemes or nonsense codons were also included. To minimize the likelihood of expressing multiple BCRs in single cells, we incorporated a T2A sequence, restricting expression to the actively expressed IGHM locus. Immediately after editing, we confirmed successful engineering by surface staining for STII, which marked 9.36% of cells (Figure S6A and left panel of Figure 4B). Genomic DNA sequencing of the edited cell population revealed that all 79 sequences were represented at high abundance (Figure S6B), confirming library integration.

To enrich BCR sequences with high surface expression, we sorted the edited OCI-Ly7 cell population into high, medium and low/negative STII staining groups (sort gating shown in the left panel of Figure 4B). Two days post-sort, we confirmed the presence of three distinct populations by repeating the surface STII staining (right panel, Figure 4B). Genomic DNA was extracted from each group, and the identity of expressed BCR sequences was determined by deep sequencing. As expected, sequences containing nonsense codons were readily detected in the low/negative STII sort group and absent from the high STII group (Figure 4C). To quantify expression, we calculated an STII enrichment score for each engineered BCR (eBCR) by comparing its relative abundance in the high STII group versus the low/negative group. These enrichment scores were used to rank each eBCR from low to high expressors (Figure S6C).

Comparison of enrichment scores between eBCRs using kappa versus lambda light chains revealed no significant differences, indicating that light chain identity does not substantially impact eBCR surface expression (Figure S6D). Lasty, we investigated whether codon optimization of human germline sequences would enhance expression. While we observed no consistent difference in enrichment scores between codon optimized (median: 0.9042) and unmodified germline sequences (median: 0.9715; Figure S6E), individual sequences showed variable enrichment, suggesting context-specific effects of optimization.

Consistent with our prior observation that AMMO1-ePCs exhibited unexpectedly low antibody secretion, the AMMO1 eBCR sequence was markedly enriched in the low-STII population (Figure 4D), indicating that BCR surface expression in the OCI-Ly7 assay may serve as a predictive marker for antibody secretion efficiency in ePCs. In contrast, several antibody sequences were highly enriched in the high STII group, including anti-SARS CoV2 clone 297^38^ (highlighted in bold in Figure S6C). To test whether BCR surface expression in the OCI-Ly7 assay correlated with secretion by ePCs, we engineered B cells to express either AMMO1 or 297 and measured antibody titers in culture supernatants. Supernatants from 297-ePCs consistently exhibited higher antibody titers than those from AMMO1-ePCs (Figure 4E). We further quantified antibody secretion at the single cell level using an anti-STII IgG ELISPOT assay, and while the number of engineered antibody-secreting cells was similar between the two groups (Figure 4F, left panel), the mean spot size was significantly larger in 297-ePCs compared to AMMO1-ePCs (Figure 4F, right panel). These results support the conclusion that BCR surface expression screening can prospectively identify antibody sequences more compatible with efficient secretion by ePCs and specifically suggest that 297-ePCs are likely to produce higher antibody titers in vivo.

### Sera from humanized mice engrafted with ePCs neutralizes SARs-CoV-2 pseudovirus

We next tested the persistence and secretion capacity of 297-ePCs in the NSG-huCD34 model. 297ePCs were administered to NSG mice bearing autologous hematopoietic grafts, as schematized in Figure 5A. In cohorts from independent experiments, we found that 297-ePCs engrafted and stably produced robust quantities of STII antibody in serum for up to 9 weeks (Figure 5B). In contrast, no STII was detected in any of the animals receiving unedited PCs. To determine whether antibody secretion in vivo was proportional to the editing rate, we measured the fraction of total serum IgG attributable to the engineered 297 antibody. Consistent with in vitro findings, this proportion closely mirrored editing efficiency across animals (Figure 5C), indicating that ePCs contributed stably to the circulating IgG pool. Importantly, in contrast to our findings with AMMO1-expressing ePCs (Figure 3G), the quantity of STII labeled antibody detected in serum was approximately 100% of the expected value based on editing rates and total IgG levels (Figure 5D). Strikingly, sera from all mice bearing 297-ePCs potently neutralized

USA-WA1/2020 SARS-CoV-2 pseudovirus (Figure 5E). Increasing serum concentrations of 297 correlated with neutralization with an approximate half maximal inhibitory concentrations of 115 ng/mL for the WA-1 receptor binding domain variant, findings similar to the published IC50 for recombinant 297 IgG.^38^ Together, these results demonstrate that human ePCs can durably engraft in an autologous humanized mouse model, produce predictable amounts of therapeutic antibody, and mediate potent antiviral activity.

## Discussion

ePCs represent an emerging cell-based modality for the sustained, high-level delivery of therapeutic proteins. Here, we describe an adoptive transfer system in which ePCs are infused into immunodeficient mice engrafted with autologous human CD34^+^ hematopoietic stem and progenitor cells. This model provides a most physiologically relevant platform that allows for robust non-clinical studies of human ePC biology, including engraftment, persistence, and delivery of therapeutic antibodies. We demonstrate that this humanized model recapitulates key features involved in PC biology, including the presence of human myeloid and T cells, as well as soluble factors such as BAFF, IL6, and IL21. Importantly, adoptive transfer of ePCs into NSG-huCD34 mice resulted in superior engraftment and durable antibody production compared to non-humanized NSG mice. We confirmed that long-lived ePCs exhibited conventional PC phenotypes in the bone marrow and spleens of NSG-huCD34 mice, and demonstrated stable, but low secretion of an anti-EBV antibody for at least 90 days. To expand the utility of this platform, we developed a BCR surface display screen to identify antibody sequences compatible with high expression and secretion. Using this approach, we selected an anti-SARS-CoV-2 antibody (clone 297) that exhibited robust surface expression in vitro, and efficient secretion by ePCs. Finally, 297-ePCs demonstrated long-term persistence and elicited potent SARS-CoV-2 pseudovirus neutralizing activity in the serum of NSG-huCD34 mice.

Human ePCs have been previously studied in immunodeficient mice, although these models have substantial limitations, including the absence of key immune niche structures and limited cross reactivity between mouse cytokines and human receptors.^19^ In prior work, we found that engraftment of ePCs into immunodeficient mice expressing human instead of mouse IL6^19^ dramatically extended ePC half-life to approximately 138 days, with circulating levels of human IgG reaching 10-100 ug/mL. An alternative strategy for supporting PCs and antibody levels in vivo involves pre-inoculation of immunodeficient mice with CD4^+^ memory T cells prior to B cell transfer.^39^ In this model, B cells differentiate in vivo in response to signals provided by the CD4^+^ T cells, which are likely activated in vivo in the presence of mouse antigens. While this approach can support IgG secretion by transferred B cells at levels far exceeding those reported in this study (>20mg/mL total huIgG^40^ and 30 ug/ml PD–1 specific immunoglobulin^13^), the kinetics differ substantially: antibody levels rise well after B cell delivery and then decline rapidly. In contrast, ePC-derived antibody levels in our model were relatively stable after infusion. The discrepancy may result from nonspecific cytokine secretion^39^ or interaction with CD4^+^ T cell transmembrane ligands that drive antigen independent polyclonal expansion of plasmablast or B cell products, some of which may subsequently differentiate into antibody secreting cells in vivo. In contrast, ePCs in NSG-huCD34 mice are exposed to physiologic levels of the PC support cytokines BAFF (10 ng/mL), IL6 (17.5 pg/mL), but not high numbers of CD4^+^ T cells (see Figure S1). We postulate that these survival signals^41^ are secreted by human support cells present in the reconstituted bone marrow and splenic niches, which likely more closely resemble the endogenous microenvironments encountered by human PCs.

While the spleen is well recognized as a niche for long-lived PCs in mice, less is known regarding its role in supporting human LLPCs. Lymphocytic choriomeningitis virus-specific PCs persist in spleen for over a year,^42^ and recent fate-mapping studies have verified the spleen as a site for PC retention.^43,44^ In our humanized model, we observed robust and sustained engraftment of ePCs in the spleens of NSG-huCD34 mice, with cells detectable up to 150 days post-infusion. This contrasts with our previous observations in human IL6 transgenic mice, where long-lived ePCs were predominantly localized to bone marrow.^19^ These data suggest that splenic persistence in the NSG-huCD34 model may be due to paracrine signaling (e.g., IL6, APRIL, BAFF) and cell-cell interactions mediated by the CD34^+^-derived human graft. Although NSG-huCD34 mice have dysfunctional myeloid cells^45,46^ and lack well-organized secondary lymphoid structures such as lymph nodes, and germinal centers, the partial reconstitution of splenic architecture that we observed here appears sufficient to support ePC survival and function. Finally, radiolabeled non-human primate PCs have been shown to efficiently engraft into both bone marrow and spleens of immune competent non-human primate recipients,^47^ reinforcing the likelihood that the spleen will serve as a key site of ePC persistence in immunocompetent human settings.

While the increased stability of antibodies produced by ePCs in serum in the NSG-huCD34 model is partly explained by support from the human cells increasing engraftment and persistence of the cells (i.e. increased bioluminescence Figure 1D), it is possible that differences in pharmacokinetics also contribute. The neonatal Fragment Crystallizable (Fc) receptor (FCRN) is critical for prolonging IgG half-life, and mouse Fcrn can cross-react with human antibodies.^48^ NSG mice exhibit decreased Fcrn expression, and accelerated clearance of IgG compared to other mouse backgrounds^49^ leading to half-life as short as 1.4 days^50^. In the NSG-huCD34 model, the presence of human myeloid cells derived from the CD34^+^ graft likely improves IgG stability through FCRN-mediated recycling, thereby enhancing antibody persistence in circulation. Therefore, the NSG-huCD34 model, through partial reconstitution of FCRN expression, may outperform NSG mice for preclinical studies of therapeutic antibody pharmacokinetics following secretion by ePCs.

We propose that key additional modifications to NSG-huCD34 mice will facilitate effective modeling of ePC-elicited antibody effector functions, including antibody-dependent cell phagocytosis, cytotoxicity, and complement lysis. For instance, binding of human IgG antibodies to Fc gamma receptors in mouse myeloid cells is similar to that in human cells,^51^ enabling the widespread use of NSG to model antibody-dependent phagocytosis^52^ that could be extended in NSG-huCD34 mice. However, NK cells, which can be supported in vivo with provision of human IL15,^53^ are required for modelling antibody-dependent cytotoxicity. Therefore, use of NSG mice expressing human IL15 for generation of NSG-huCD34 mice could, support development of functional natural killer cells from human CD34 grafts,^53,54^ and facilitate modeling of antibody-dependent cellular cytotoxicity via NK cells.^55^ While NSG mice lack the hemolytic complement gene,^56^ an NSG model^57^ with the hemolytic complement gene has been created that could be used for evaluation of IgM-mediated complement lysis driven from IgM antibodies secreted by ePCs. Finally, evaluation of human ePCs in near-immunocompetent settings may be possible using KIT mutant mice humanized with cord blood and treated with estrogen.^58^ These mice develop functional immune follicles across several tissues,^59^ and future studies should prioritize development of engineering protocols for cord blood derived B cells to enable testing of autologous ePCs in this setting.

Similar to prior reports,^10,12^ we observed variable protein expression from different antibody sequences in both a human B cell line and in primary human PCs. This variability is likely driven by sequence-dependent effects on translation efficiency, post-translational modifications, and folding. Incomplete formation of disulfide bonds during antibody folding^60^ can lead to retention of antibodies in the endoplasmic reticulum^61^, degradation of the misfolded antibodies, and activation of the unfolded protein response. Additionally, non-canonical cysteine^62^ or hydrophobic^63^ residues with the complementary determining regions in antibodies have been associated with aggregation and impaired secretion across multiple systems. Codon usage may also contribute as B cells and PCs exhibit codon usage patterns distinct from other cell types,^64^ and suboptimal codon composition could negatively impact translation efficiency.

Because many of the antibody sequences in our libraries were identified from memory B cells using antigen-specific tetramers,^38^ it is not surprising that they may not be optimized for the robust expression and secretion in ePCs. These findings underscore the importance of selecting antibody sequences not only for antigen specificity, but also for compatibility with the secretory machinery of plasma cells, a process that could be facilitated using our surface display method.

In summary, we have generated a robust humanized mouse model for ePC adoptive transfer that recapitulates many key aspects of human PC biology and enables functional evaluation of therapeutic antibody delivery. This platform not only provides a tool for studying fundamental PC behavior in vivo but also offers a powerful preclinical system for the development and testing of ePC-based cell therapies.

## Methods

### Generation and Analysis of NSG-huCD34 mice

All animal studies were performed according to AAALAC standards and were approved by the Seattle Children’s Research Institute (SCRI) Institutional Animal Care and Use Committee. NOD.Cg-Prkdcscid Il2rgtm1Wjl/SzJ-c (NSG) mice were purchased from Jackson Laboratory and all mice were kept in a designated pathogen-free facility at SCRI. A general list of reagents used for cell culture and processing is described in Table S1. Human peripheral blood stem cells were obtained from healthy G-CSF mobilized donors in accordance with Seattle Children’s Research Institute’s institutional review board. Frozen human CD34^+^ cells were thawed and then cultured in SCGM (CellGenix) with 100 ng/mL each of human recombinant thrombopoietin, stem cell factor, and Flt3-ligand at a cell density of 1×10^6^ cells/mL in a humidified 37°C incubator with 5% CO2. After 24 hours in culture, cells were washed in phosphate buffered saline (PBS) and 0.5-1×10^6^ CD34^+^ cells delivered per busulfan-conditioned NSG mouse by retro-orbital injection. Busulfan conditioning was performed by intraperitoneal (i.p.) injection of 35 mg/kg clinical grade busulfan into 7-8 week-old NSG mice 24 hours prior to cell transfer. 6-10 weeks after transfer, successful human cell engraftment was confirmed by the presence of human CD45+ cells in peripheral blood using flow cytometry. 10 weeks after transfer, NSG-huCD34 and matched NSG littermates were harvested for peripheral tissues. Half of the spleens were fixed in 10% neutral pH formalin and sent to the Fred Hutch Experimental Histopathology for H&E staining. Half of the spleens were processed and analyzed by flow cytometry. Peripheral blood was processed for sera and then cytokine concentrations were measured via multiplexed ELISA (U-PLEX, Meso Scale Discovery).

### Homology directed repair of B cells using CRISPR Cas9 and AAV6 repair templates

Mobilized PBMCs were collected from the left over CD34^-^ fraction from mobilized donors in accordance with Seattle Children’s Research Institute’s institutional review board. We isolated B cells using the EasySep Human B cell Enrichment Kit (Stem Cell Technologies). Isolated B cells were cultured in Iscove’s modified Dulbecco’s medium (Gibco), supplemented with 2-mercaptoethanol (55μM) and 10% FBS and human cytokines. Cells were cultured in activating and differentiating conditions as described in Cheng et al (Figure 2A).^65^ Cells for *in vivo* experiments were purified via CD3 bead depletion column (Miltenyi) immediately prior to tail vein injection. Cells were phenotyped by flow cytometry.

### Generation of trackable ePCs via AAV6-mediated gene editing

B cells and subsequent PCs were engineered to express the trackable reporter *firefly* luciferase or BFP as previously described in Cheng et al.^19^ Digital droplet PCR was performed to calculate the HDR allele frequency as previously described in Hung et al.^22^ BFP expression was confirmed by flow cytometry.

### Tracking and phenotyping of ePCs in NSG-huCD34 mice

One to ten million luciferase or BFP-expressing ePCs were injected intravenously into NSG and NSG-huCD34 mice. ePC engraftment was monitored by bioluminescence imaging using IVIS Lumina S5 (Perkin Elmer) following subcutaneous injection of luciferin (75-150 mg/kg). Peripheral blood was collected by retro-orbital bleed or submandibular bleed and processed to collect sera and quantify peripheral IgG titers (Invitrogen human total IgG ELISA Kit). A subset of the mice were euthanized for ex vivo spleen and bone marrow immunophenotyping by flow cytometry. See Table S2 for all flow antibodies used in the study.

From some mice, tissues were fixed in 10% neutral pH formalin and provided to Fred Hutch Experimental Histopathology for immunohistochemical analysis.

### BCR Surface screen generation and analysis

OCI-Ly7 cell line (DSMZ) was cultured in IMDM (Gibco) media supplemented with 10% FBS. Clustered regularly interspaced short palindromic repeats (CRISPR) RNAs (crRNAs) targeting the second exon of IGHM were identified using the Broad Institute GPP sgRNA (Table S3) Designer and synthesized (IDT) containing phosphorothioate linkages and 2′O-methyl modifications. crRNA and tracrRNA; single guide hybrids were mixed with 3uM Cas9 nuclease (Berkeley Labs) at a 1.2:1 ratio and delivered to cells by Lonza 3D (CM-137) electroporation.

After electroporation, cells were incubated in the presence of adeno-associated virus 6 (AAV6) vectors carrying homologous DNA repair templates consisting of a T2A ribosomal skip element in cis with a light chain linked via a 3x strep tag II (STII) sequence to the variable domain of a heavy chain linked to a codon optimized version of the first exon of IGHM flanked by 400bp homology (20% AAV). We developed a gibson assembly-based drop in cloning method to rapidly generate a library of gene pAAV transfer plasmids containing 79 unique eBCR sequences as well as additional 15 codon variations. The majority of the pool of sequences consisted of antibodies targeting the receptor binding domain of SARS-CoV2 spike protein (14 clinical monoclonal antibodies, 29 published^38,66,67^, 14 unpublished). An additional 10 known control eBCRs that do not bind receptor binding domain (ie Palivizumab, VRC01, MSP1 etc), 6 nonsense controls and 10 eBCR sequences that were codon optimized (codon optimization tool, IDT) away from germline sequence were included in the library. Another 5 of the eBCR sequences had their signal peptide swapped to each of three VH signal peptides for an additional 15 sequences. Each eBCR sequence had its IgHM exon 1 silently mutated to a unique barcode sequence to allow for easy sequencing and tracking of each sequence. These barcodes were amplified by nested PCR and sequenced using a miseq nano kit v3. Barcodes were counted using 2Fast2Q.^68^ Each eBCR count was normalized to total counts from each sample to get sequence frequency. OCI-L7 cells expressing the eBCR library were sorted via FACS ARIA II based on staining of the STII linker. By comparing sequence frequency in the high STII group to the low STII group we were able to calculate an enrichment score.

### Generation of AMMO1- and 297-secreting ePCs and detection of STII-IgG and human total IgG

B cells were engineered to express an engineered version of the anti-EBV antibody, AMMO1^35^, or anti-SARS-CoV-2 antibody, 297^38^, as previously described in Moffett et al.^12^ Engineered B cells were differentiated to PCs as previously described in Cheng et al.^65^ Editing rates were calculated by intracellular Strep Tag II staining of AMMO1 ePCS. AMMO1 was detected via ELISA using a gHgL-based capture reagent (provided by Andy Mcguire) and human peroxidase conjugated anti-human IgG detection antibody (Jackson). Anti-SARS-CoV-2 antibody, 297, was detected using anti-NWSHPQFEK (STII) capture antibody (Genscript) and peroxidase anti-human IgG detection antibody (Jackson). Total human IgG and total human IgM was determined using IgG (total) human ELISA kit (Invitrogen). The expected AMMO1 and 297 titers were determined by multiplying the STII^+^ cell frequency by the total IgG found in the sera or supernatant. Additionally, ePCs were assessed for antibody secretion by STII enzyme-linked immunospot (ELISPOT) assay. Briefly, various dilutions of cells were seeded in an anti-STII coated (Genscript) Millipore plate at and incubated overnight at 37 °C in 5% CO2. After 24 hours, plates were washed with 1x PBS containing 0.05% Tween 20, before subsequent incubation with peroxidase conjugated goat anti-human IgG (Southern Biotech). Then, plates were washed. The plates were washed four times and developed using AEC Chromogen (BD Biosciences, Canada). The plates were then rinsed with distilled water and dried at room temperature overnight. Results were obtained and analyzed using IRIS plate imager (MabTech).

### Neutralization of SARS-CoV-2 pseudovirus

Pseudovirus neutralization assays were performed as previously described.^38,67,69^ Briefly, ACE-2–expressing 293T cells (BEI Resources; NR-52511) were seeded onto poly-L-lysine– coated 96-well plates and grown to 85–95% confluency. WA-1 SARS-CoV-2 pseudotyped lentivirus was incubated with serially diluted sera from ePC engrafted mice or 297 recombinant antibody for 1 h at 37°C and then gently applied to cells. 48 h after infection, cells were lysed following the manufacturer’s instructions using the Steady-Glo Luciferase Assay System reagent (Promega), and luminescence was measured in black-bottom plates using a Centro LB Microplate Luminometer (Berthold Technologies) with MikroWin 2000 software set to a 1-s exposure time. Luminescence values were normalized relative to the luminescence in control wells that had been transduced with virus coincubated with control NSG sera (internal for each plate, average of six wells).

### Statistical analysis

Statistical analyses using parametric tests were performed using Prism 10 (GraphPad, San Diego, CA) as described in figure legends.

## Acknowledgements

The authors graciously thank Gene Hess for technical flow cytometry assistance, the Office of Animal Care for technical husbandry training and assistance, and the SCRI viral core for making AAV6 virus.

This research was supported by National Institutes of Health (NIH) grants F30 5F30AI164574-02 (T.F.H.; National Institute of Allergy and Infectious Disease [NIAID]). This work was also supported in part by the Seattle Children’s Research Institute (SCRI) Program for Cell and Gene Therapy (PCGT), the Children’s Guild Association Endowed Chair in Pediatric Immunology (to D.J.R.), and the Hansen Investigator in Pediatric Innovation Endowment (to D.J.R.). The work was also supported by the National Cancer Institute under activity number 5R01CA201135 (to R.G.J.). Finally, the work was partially supported by a research agreement with Be Biopharma. These funding sources had no influence on the design of the study; collection, management, analysis, and the interpretation of the data; preparation, review, or approval of the manuscript; and the decision to submit the manuscript for publication.

## Authorship Contributions

Contributions: TFH, PN, MH, NDC, CDH, SL and ATM designed and constructed the antibody sequences, homology-directed repair templates and AAV used in the study. TFH, LH, and ATM evaluated the in vitro function of the AMM01 antibody and sera from mice. TFH, AEH, RYHC, ERS, ARO, and CMS did the in vitro studies with human B cells and ePCs. TFH, AEH, SS, GDA, and CMS conducted the in vivo studies. TFH developed the surface display screen and conducted the immunohistochemistry and secreted proteins studies. TFH, RGJ and DJR designed the study. TFH, AEH, RGJ and DJR wrote and edited the manuscript.

---

**Table S1.** Reagents.

| Item | Vendor | Catalog No. |
| --- | --- | --- |
| OCI-Ly7 | DSMZ | ACC 688 |
| ACE-2 expressing 293Ts | Provided by Bloom lab |  |
| IgG (total) Human Uncoated ELISA kit | Invitrogen | 50-112-8849 |
| Peroxidase AffiniPure Goat Anti-Human IgG | Jackson ImmunoResearch | 109-035-003 |
| Goat Anti-Human IgG-HRP | Southern Biotech | 2040-05 |
| NWSHPQFEK antibody, pAb, Rabbit (anti-STII linker) | GenScript | A00626 |
| NWSHPQFEK Tag Antibody (HRP), mAb, Mouse | GenScript | A01742 |
| AEC Substrate Kit, Peroxidase (HRP), (3-amino-9-ethylcarbazole) | Vector Laboratories | SK4200 |
| EasySep HumanB cell Enrichment kit | StemCell Technologies | 19054 |
| CD3 Microbeads, human, depletion kit | Miltenyi Biotec | 130-097-043 |
| SteadyGlow | Promega | E2510 |
| SCGM | CellGenix | 20806-0500 |
| thrombopoietin | Peprotech | 300-18 |
| stem cell factor | Peprotech | 300-07 |
| Flt3-ligand | Peprotech | 300-19 |
| DMEM | Corning | 15-013-CV |
| GlutaMAX | Gibco | 35050061 |
| IMDM | Gibco | 12-440-053 |
| 2-mercaptoethanol | Gibco | 21985023 |
| Fetal Bovine Serum | Sigma | 12103C |
| Multimeric CD40L | Adaptogen | AG-40B-0010 |
| CpG oligodeoxynucleotide 2006 | Invitrogen | A10168 |
| IL-6 | PeproTech | 200-06 |
| IL-15 | Miltenyi | 130-095-765 |
| IL-21 | PeproTech | 200-21 |
| IFN alpha 2B | Proteintech | HZ-1072 |
| Cas9-NLS purified protein | UC Berkeley MacroLab |  |
| tracrRNA | IDT | 1072534 |
| P3 Primary Cell 4D Nucleofector X kit S | Lonza | V4XP-3032 |
| MiSeq Reagent Kit v3 | Illumina | MS-102-3001 |
| NOD.Cg-Prkdcscid Il2rgtm1Wjl/SzJ-c (NSG) mice | Jackson Laboratory | 005557 |
| Busulfan | Selleckchem | S1692 |
| Pierce D-Luciferin Monosodium Salt | Thermo Fisher | 88292 |
| Paraformaldehyde | Thermo Fisher | AA47377-9M |

**Table S2.** Flow Cytometry reagents.

| Target Antigen | Clone | Fluorophore | Dilution | Vendor | Catalog Number |
| --- | --- | --- | --- | --- | --- |
| Live/Dead |  | AF350 | 1:1000 | Thermo Fisher | A10168 |
| Human CD3 | OKT3 | BV605 | 1:300 | BioLegend | 317322 |
| Human CD4 | OKT4 | BV650 | 1:300 | BioLegend | 317436 |
| Human CD19 | HIB19 | PE-Cy7 | 1:300 | eBioscience | 25-0199-42 |
| Human CD33 | WM53 | AF700 | 1:300 | BioLegend | 303436 |
| Human CD38 | HIT2 | PerCP-Cy5.5 | 1:300 | BD | 551400 |
|  |  |  |  | Biosciences |  |
| Human CD45 | 2D1 | APC-Cy7 | 1:300 | BioLegend | 368516 |
| Human CD138 | DI101 | PE | 1:50 | BioLegend | 352306 |
| Human IgM | MHM-88 | APC | 1:300 | BioLegend | 314510 |
| Human IgG | G18-145 | AF700 | 1:300 | BD Biosciences | 561296 |
| NWSHPQFE K (STII) | 5A9F9 | FITC | 1:500 | Genscript | A01736 |
| Murine CD45 | 30-F11 | BV421 | 1:500 | BioLegend | 103133 |

**Table S3.**
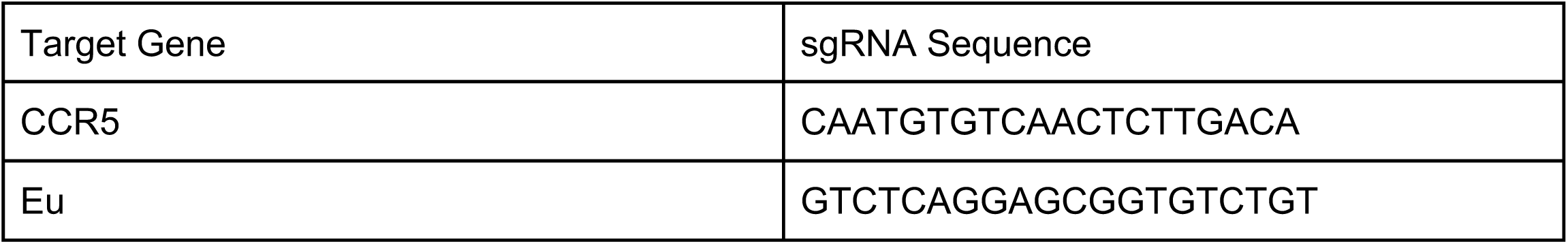
Guide RNAs.

**Table S4.**
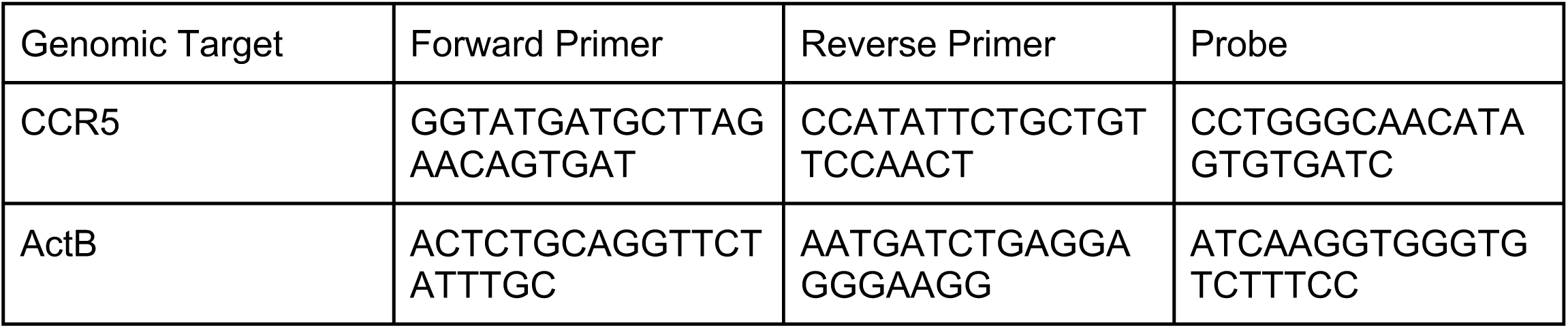
ddPCR reagents.

**Figure S1:**
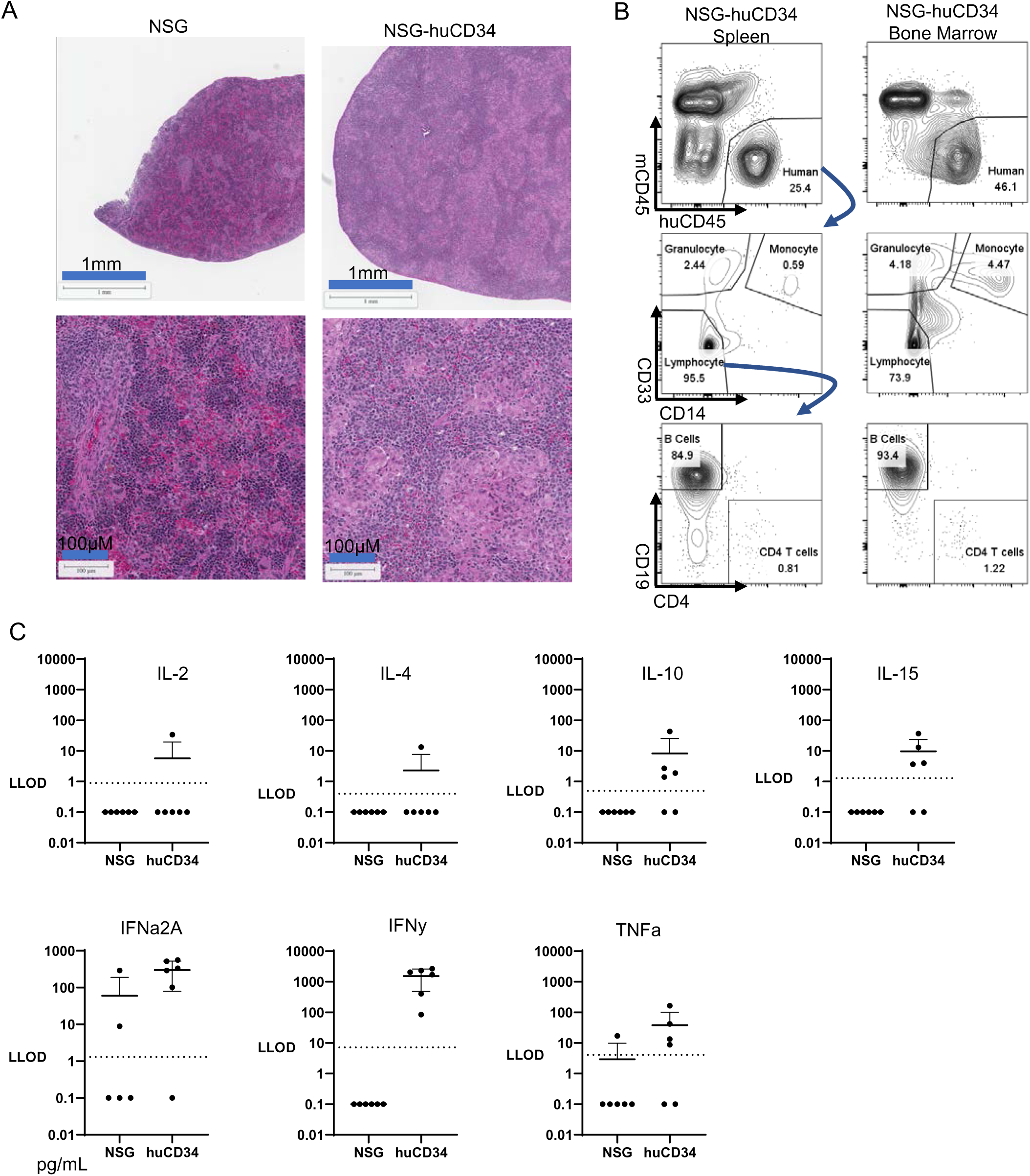
Humanization of NSG mice with human CD34+ cells partially reconstitutes human PC niches. Immunodeficient NSG mice were injected with huCD34^+^ hematopoietic stem and progenitor cells via tail vein injection. Mouse tissues and peripheral sera were harvested 10 weeks after engraftment. **A)** Representative H&E images of spleens from NSG and NSG-huCD34^+^ mice shown. **B)** Representative flow plots of cells harvested from a NSG-huCD34 mouse spleen and femur bone marrow that were processed, stained for human surface markers. Blue arrows indicate flow gating of different cell populations. **C)** Cytokine concentrations in the peripheral sera of NSG-huCD34 and NSG mice. LLOD stands for lower limit of detection. Data from two donors in two independent experiments.

**Figure S2:**
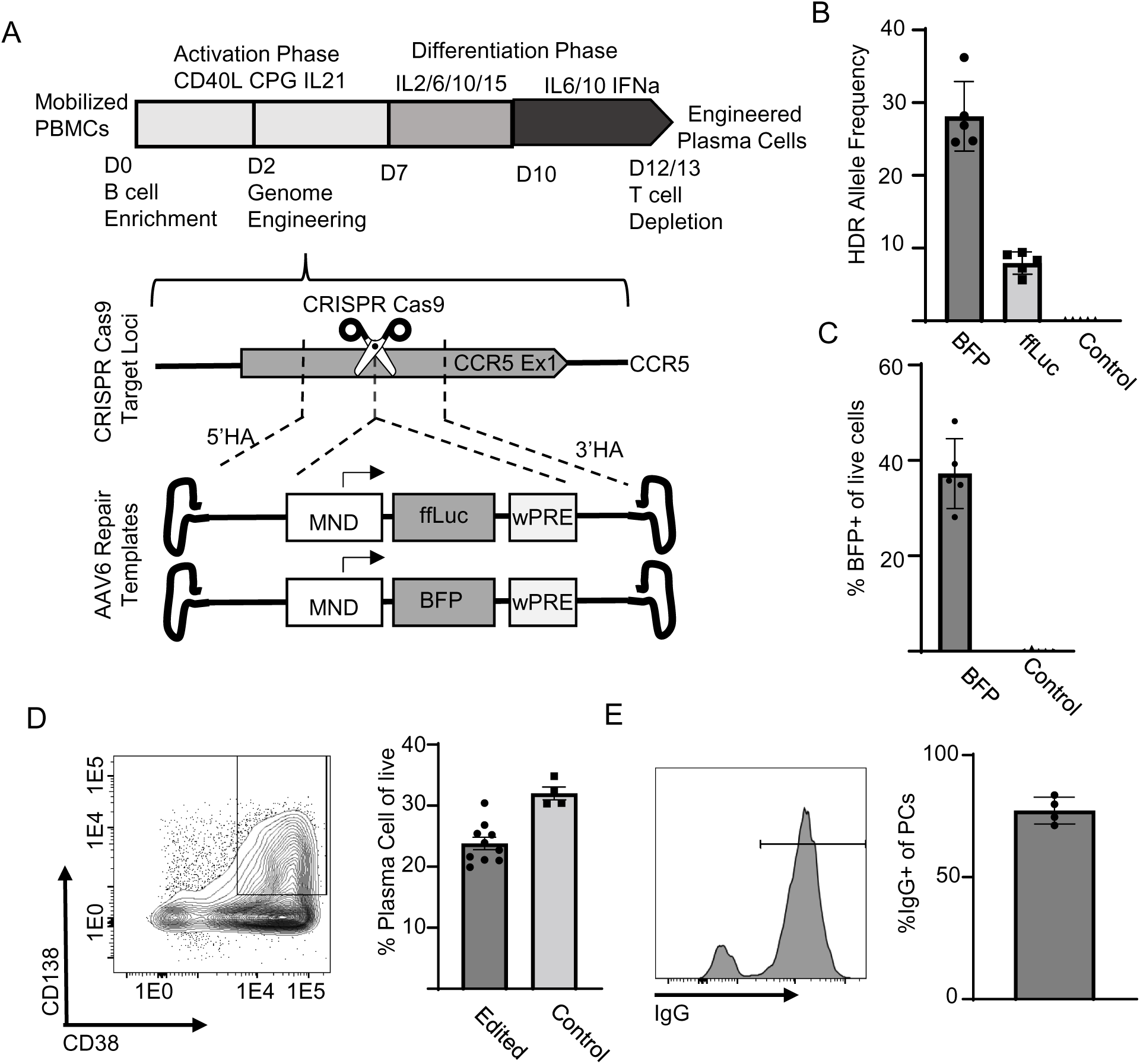
Genome-engineering of mobilized primary human B cells followed by differentiation generates trackable ePCs. **A)** Schematic indicating the *in vitro* culturing and genome engineering strategies to generate trackable ePCs from mobilized PBMCs. **B)** HDR allele frequency as calculated by ddPCR (Table S4) of 40ng of genomic DNA from ePCs. **C)** The percent BFP^+^ of day 13 ePCs as measured by flow cytometry. **D)** Representative flow cytometry plot of ePCs (gated on live, single cells) at the end of the differentiation phase. Gate shown for the quantification of cells expressing both PC markers CD38 and CD138 shown in the right bar graph. Intracellular staining IgG was performed on D13 ePCs. **E)** Representative flow plot and quantified percent IgG^+^ intracellular staining of live CD38^+^ CD138^+^ PCs. Data from 5 donors across three independent experiments.

**Figure S3:**
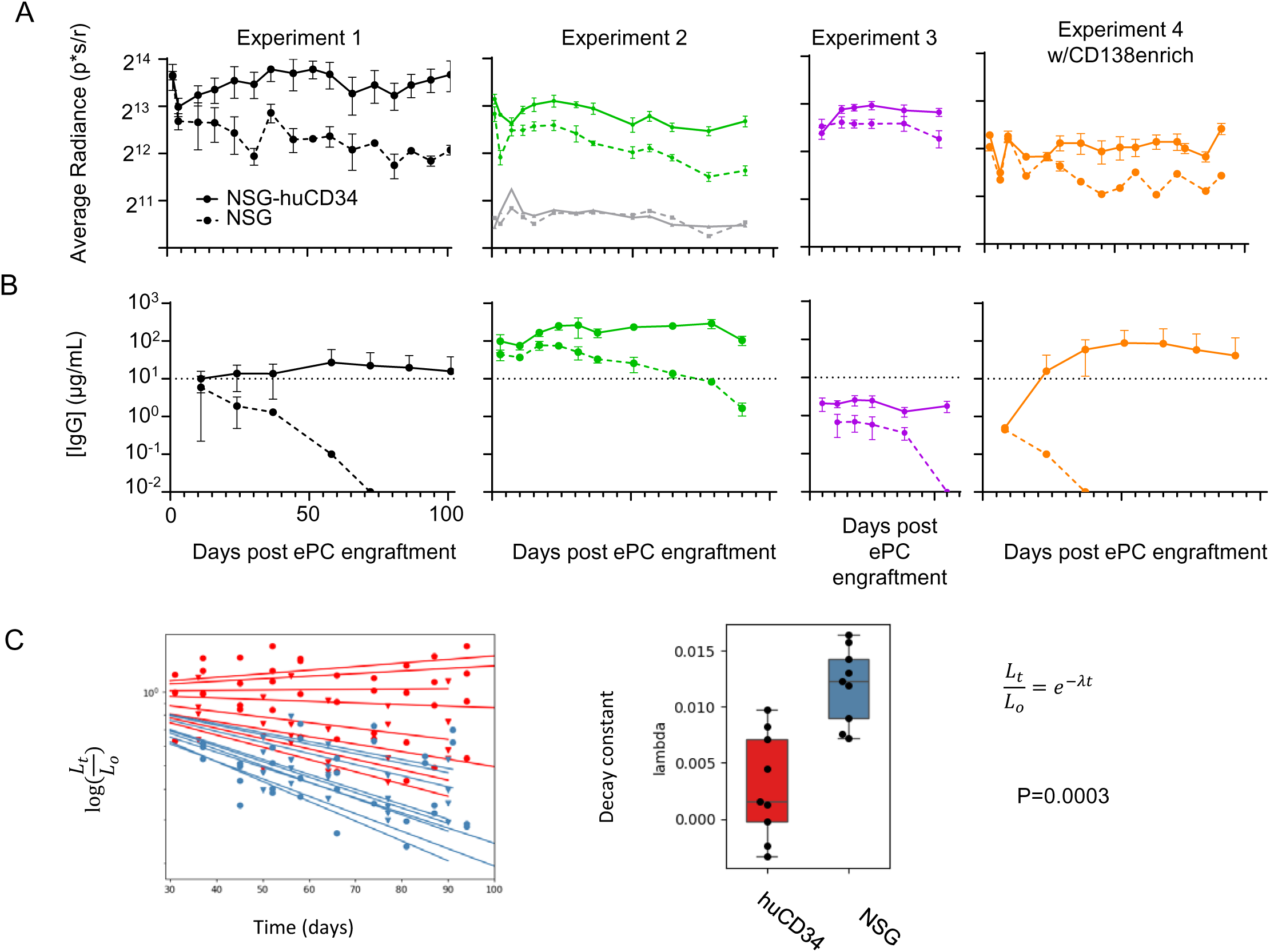
ePCs have superior engraftment in humanized mice across multiple experiments. Immunodeficient NSG mice were injected with huCD34^+^ hematopoietic stem and progenitor cells or PBS via tail vein injection. Ten weeks later, NSG and NSG-huCD34 mice received luciferase expressing ePCs. Mice were subcutaneously injected with luciferin and imaged at various time points. **A)** Average radiance was measured, averaged across experimental groups and then plotted by experiment. **B)** The concentration of human IgG was measured via ELISA of peripheral sera averaged by experimental group and plotted for each independent experiment. A dotted line was included at 10 μg/mL to aid in the assessment of the data. **C)** For each animal with greater than 5 time points data beyond day 30 (NSG, n = 9 animals; NSG-huCD34, n = 10 animals), we used the fitted curves to calculate the decay rate based on the bioluminescence data in Figure 1D. The decay rates were then used to calculate antibody half-life. A paired student’s t-test was used to calculate p-values for the decay constants (p = .0003).

**Figure S4:**
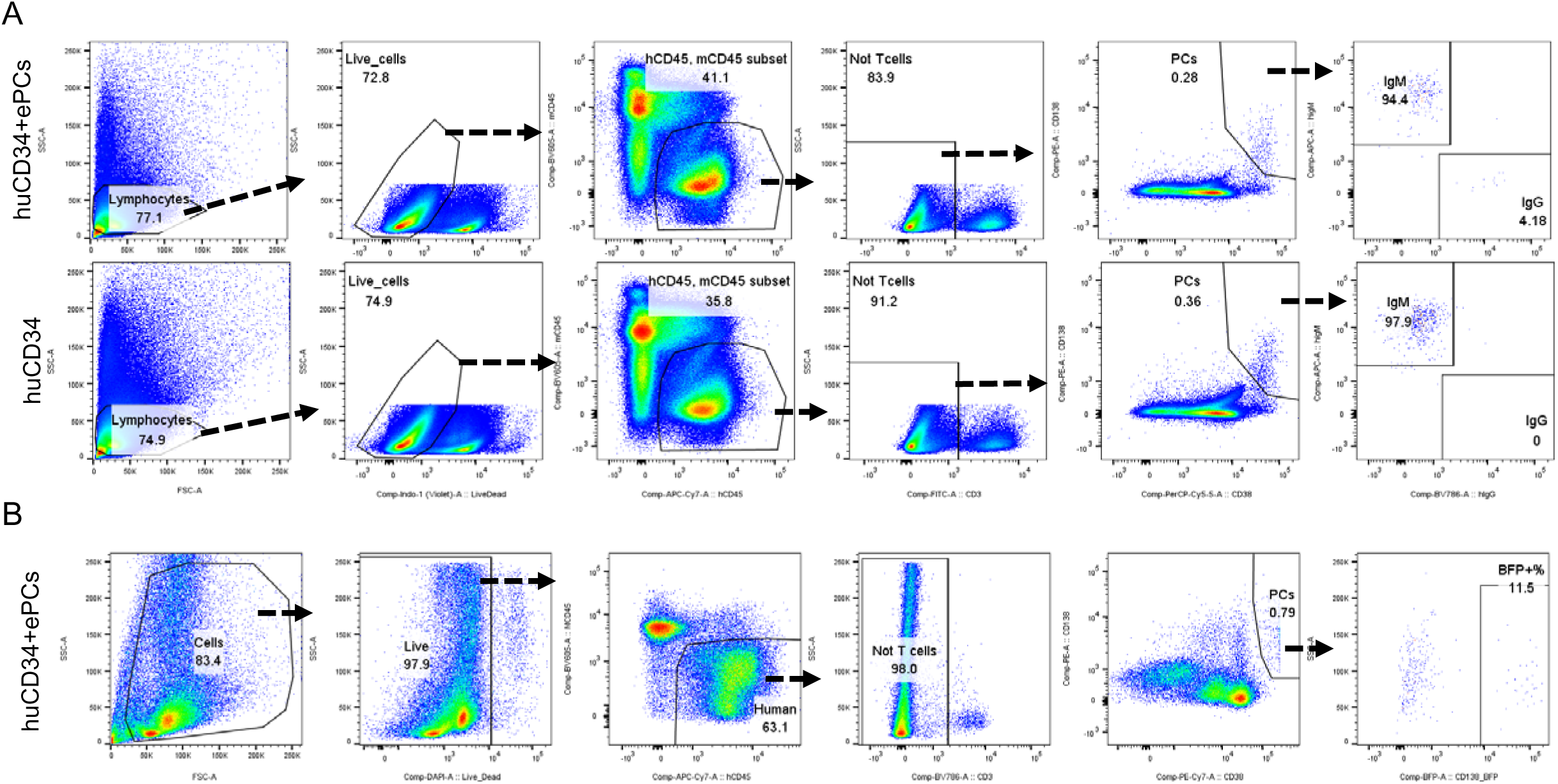
Phenotype and location of ePCs recovered from tissues of NSG-huCD34 mice. Mouse bone marrow and spleen were harvested between 5 and 10 weeks post ePC engraftment from NSG or NSG-huCD34 mice. **A)** Representative flow gating strategy used to determine the isotype of ePCs recovered from the spleen and bone marrow. **B)** Representative flow gating strategy used to determine the percent of ePCs that express BFP. **C)** Representative immunohistology images of bone marrow (vertebrae) and spleen from NSG-huCD34 mice engrafted with luciferase expressing ePCs.

**Figure S5:**
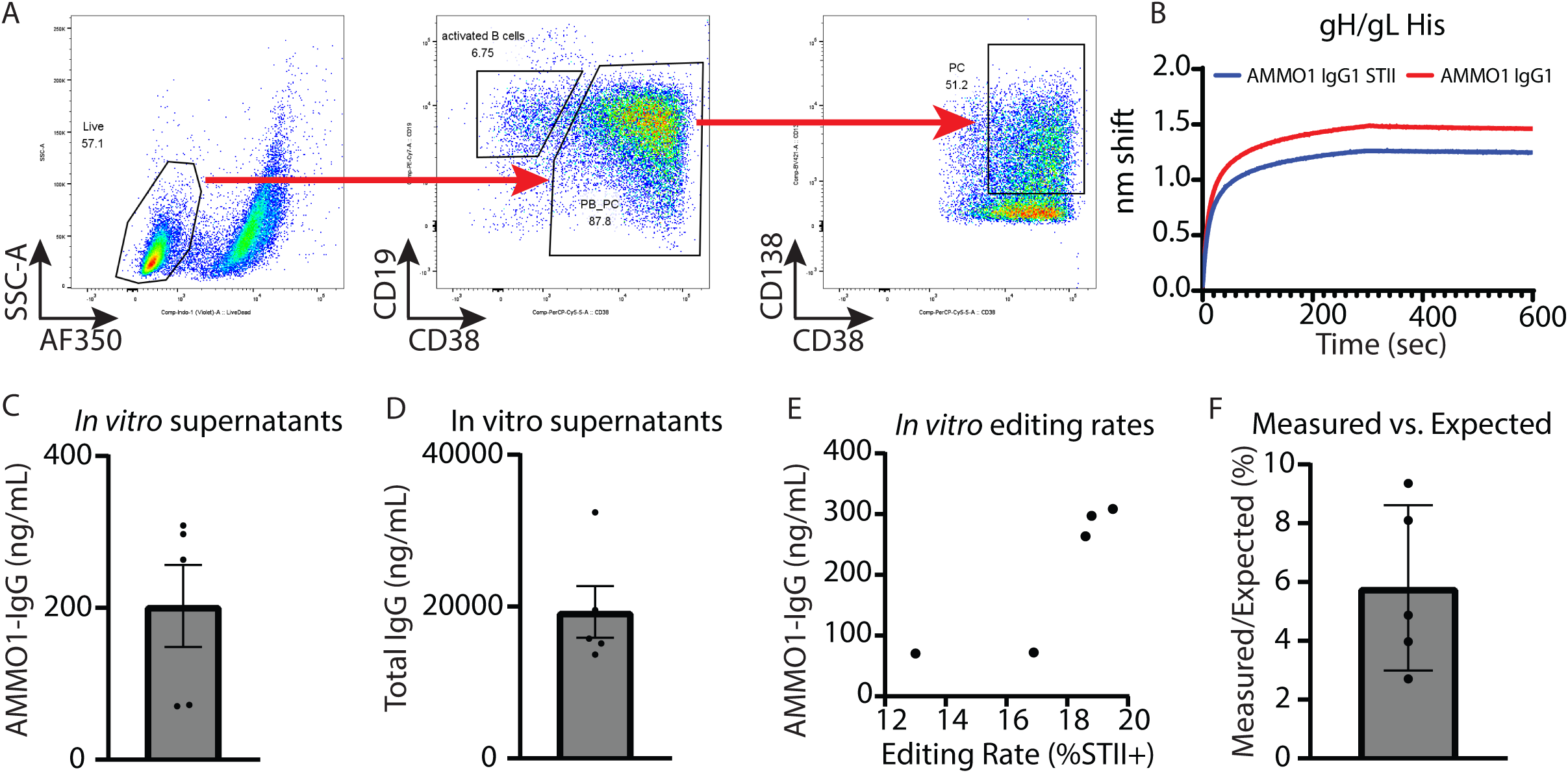
In vitro characteristics of AMMO1 ePCs. **A)** Representative flow cytometry gating of *in vitro* differentiated PCs. **B)** AMMO1-IgG1 with separate heavy and light chains, along with AMMO1 expressed as a single chain STII fused IgG1 were produced from HEK 293T cells and purified. The fused and split AMMO1 antibodies were assessed for affinity via gHgL biolayer interferometry. **C)** AMMO1-IgG levels as measured by gHgL ELISA from day 10 supernatants of AMMO1 expressing ePCs. **D)** Total human IgG levels as measured by total human IgG ELISA from day 10 supernatants of AMMO1 expressing ePCs. **E)** Supernatant AMMO1-IgG levels were compared to editing rates (% STII^+^ by flow cytometry). **F)** We compared the measured AMMO1 concentration to the expected AMMO1 concentration based on AMMO1 editing rates measured by flow cytometry and graphed the ratio.

**Figure S6.**
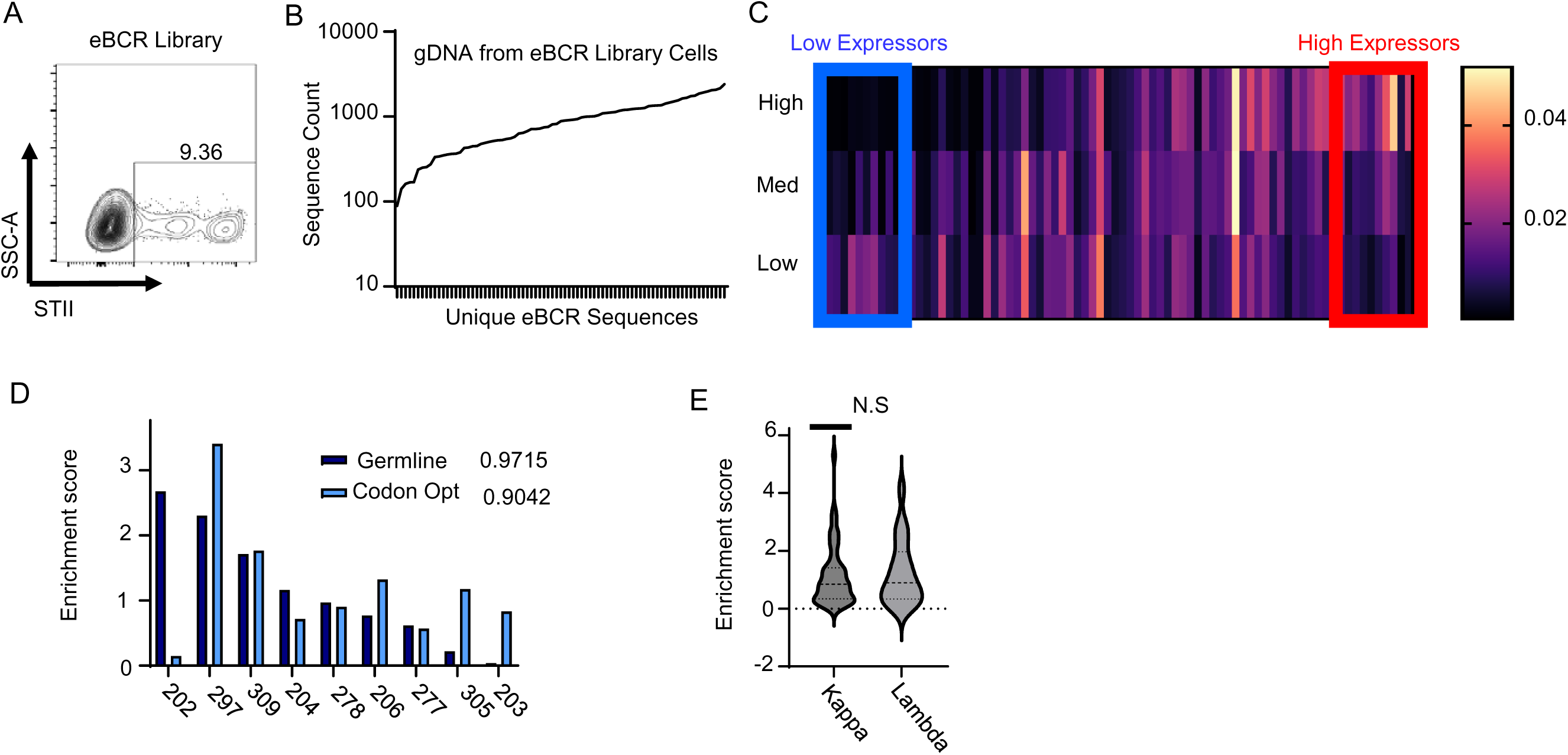
BCR surface display screen. A) OCI-Ly7, a diffuse large B-cell lymphoma cell line, was engineered to express a library consisting of 79 different BCR sequences engineered with an STII linker between the heavy and light chains. BCR library engineered OCI-Ly7 were surface stained for STII. B) Genomic DNA from the bulk BCR library engineered OCI-Ly7 was collected and deep sequenced. C) Enrichment scores were calculated by comparing the sequence prevalence of each eBCR in the high STII sorted group to the low STII sorted group. Sequence prevalence for each eBCR was plotted from low to high STII enrichment score. The blue box indicates “low expression” eBCR sequences that have low STII enrichment scores while the red box indicates “high expression” sequences with high STII enrichment scores. D) Enrichment score of eBCRs expressed as germline sequence or as codon optimized sequence. Each BCR sequence was identified and graphed according to sequence count. E) Violin plot showing the enrichment score of eBCRs with kappa light chains or lambda light chains.

